# Early life adversity decreases fear expression in pre-adolescence by accelerating amygdalar parvalbumin cell development

**DOI:** 10.1101/2020.01.07.892687

**Authors:** Gabriela Manzano Nieves, Marylin Bravo, Kevin G. Bath

## Abstract

Resource insecurity (e.g., poverty) can be a significant source of stress. Decreased resources during childhood has been associated with increased risk for developing stress-related disorders, including major depressive disorder and anxiety Although the link between early life adversity and increased risk for psychopathology has been well established, the developmental mechanisms remain unclear. Using a mouse model of poverty-like rearing, limited bedding and nesting materials (LB), we tested the effects of LB on the development of fear learning and of key neuronal structures involved in emotional regulation, the medial prefrontal cortex (mPFC) and basolateral amygdala (BLA). LB delayed the ability of pre-adolescent mice to express, but not form, an auditory conditioned fear memory. LB disrupted typical fear circuit development, accelerating parvalbumin positive (PV+) inhibitory interneuron maturation in the BLA and delaying the maturation of connections between the mPFC and BLA. The decreased fear expression in LB reared mice during early development was rescued through optogenetic inactivation of PV+ cells in the BLA. Together our data demonstrate that LB has profound and deleterious effects on mPFC and BLA development, decreasing threat-associated behavior expression, but not learning, in childhood. The current results provide a model of transiently blunt emotional reactivity in childhood, with fear-associated memories emerging later in adolescence, and possibly contributing to later pathology development.

## Introduction

Early life adversity (ELA) increases the lifetime risk for multiple forms of psychopathology, including anxiety disorders and major depressive disorder (Agid et al., 1999; Draijer and Langeland, 1999; Widom, 1999; Heim and Nemeroff, 2001; Koenen and Widom, 2009). ELA is associated with higher rates of negative outcomes than similar events experienced in adulthood (Mullen et al., 1996; McCauley et al., 1997; Felitti et al., 1998; Heim and Nemeroff, 2001; Salmon and Bryant, 2002). The increased lifetime risk for psychopathology is proposed to result from alterations in the developmental trajectories of brain centers regulating emotional learning and emotional expression. In agreement with this prediction, ELA has been shown to drive early engagement of the basolateral amygdala (BLA), a key node supporting emotional processing, threat assessment, and fear learning (Moriceau et al., 2009; Bath et al., 2016).

The neural circuit(s) supporting fear and aversive learning in adult rodents have been well characterized. Numerous studies have shown that the prelimbic (PL) subregion of the medial prefrontal cortex (mPFC) projects to the BLA (Vertes, 2004; Do-Monte et al., 2015b) and is necessary for fear retrieval (Burgos-Robles et al., 2009; Sierra-Mercado et al., 2011; Courtin et al., 2014), whereas the infralimbic (IL) subregion of mPFC supports extinction learning (Sierra-Mercado et al., 2011; Adhikari et al., 2015; Do-Monte et al., 2015a). Less is known about the development of fear learning circuits. PL projections into amygdala begin to emerge around postnatal day (PND) 7, increasing through adolescence (Cunningham et al., 2002), and being pruned back during early adulthood (Bouwmeester et al., 2002; Cressman et al., 2010). The relative late integration of the PL into the threat learning circuit may explain recent reports suggest that the infant PL, unlike adult PL, is not involved in sustaining fear responses (Chan et al., 2011). Understanding how the fear circuit is developing can help us predict differences in the acquisition and expression of learned aversive behaviors.

Recently, a rodent model of resource insecurity, limited bedding and nesting (LB) was developed to simulate aspects of poverty (Rice et al., 2008; Bolton et al., 2019). LB has been shown to induce significant distress in the dam, alter patterns of maternal behavior (McLoyd, 1998; Rice et al., 2008; Bolton et al., 2019), and delay developmental processes such as sexual maturation and physical growth (Yam et al., 2017; Manzano Nieves et al., 2019). In addition, LB impacts behavioral outcomes with effects that persist into adulthood, increasing depressive-like behaviors (Goodwill et al., 2019), altering behavioral response to stress (Cohen et al., 2013; Manzano-Nieves et al., 2018), and affecting hippocampal dependent learning (Wang et al., 2011; Bath et al., 2016, 2017; Manzano-Nieves et al., 2018). However, the effects of LB on the development of neuronal populations and regions supporting fear learning and expression remain unclear. Here, we tested the impact of LB on the development of subregions of mPFC, BLA, projections from mPFC to BLA, as well as the effect on fear learning and expression.

Here we show that LB delayed somatic development in mice, altered developmental expression of fear learning, and had adverse effects on the development and connectivity of mPFC and BLA. Specifically, LB resulted in decreased body weight and brain weight from infancy into pre-adolescence. Changes in physical development were accompanied by an acceleration in parvalbumin cell differentiation in the BLA of pre-adolescent mice and decreased mPFC to BLA anatomical connectivity during adolescence. Furthermore, LB mice exhibited decreased fear expression during pre-adolescence, which was rescued through optogenetic inactivation of parvalbumin cells in the BLA. Based upon these results premature differentiation of parvalbumin cells in BLA following poverty-like rearing leads to a transient decrease in fear expression.

## Results

### Early life adversity decreases physical growth

In humans, early life adversity is correlated with changes in expected weight (Rondó et al., 2003; Hult et al., 2010; Wainstock et al., 2013; Maniam et al., 2014), with low weight in infancy predicting later cognitive deficits (Strathearn et al., 2001; Corbett and Drewett, 2004). Thus, weight gain may serve as a biomarker for altered neurodevelopment and later risk for pathology. To model a poverty-like conditions, dams and pups were placed in conditions of low bedding and nesting materials from PND 4–11 (McLoyd, 1998; Rice et al., 2008; Bolton et al., 2019). To assess the impact of LB on somatic and brain development, the body weight and brain weight of control (ctrl) and LB reared mice was measured at select ages across early development (PND 16, 21, 28, and 35). LB mice weighed significantly less than control mice at PND 16, 21, and 28, with differences diminishing by PND 35 (**Figure 1B left**). Examination of brain weight (**Figure 1B center**) revealed a similar pattern, with the brains of LB mice weighing less than control brains at PND 16, 21, and 28, but not at PND 35. Early adversity may shift resources away from somatic development to support brain development. To test this prediction, the brain to body weight ratio was calculated across developmental time points (**Figure 1C right**). LB reared mice had a higher brain-to-body weight ratio than control mice at PND 21 and PND 28. Thus LB effects on brain and total body weight were not proportional, with a greater impact on body development at select developmental time points.

**Figure 1.**
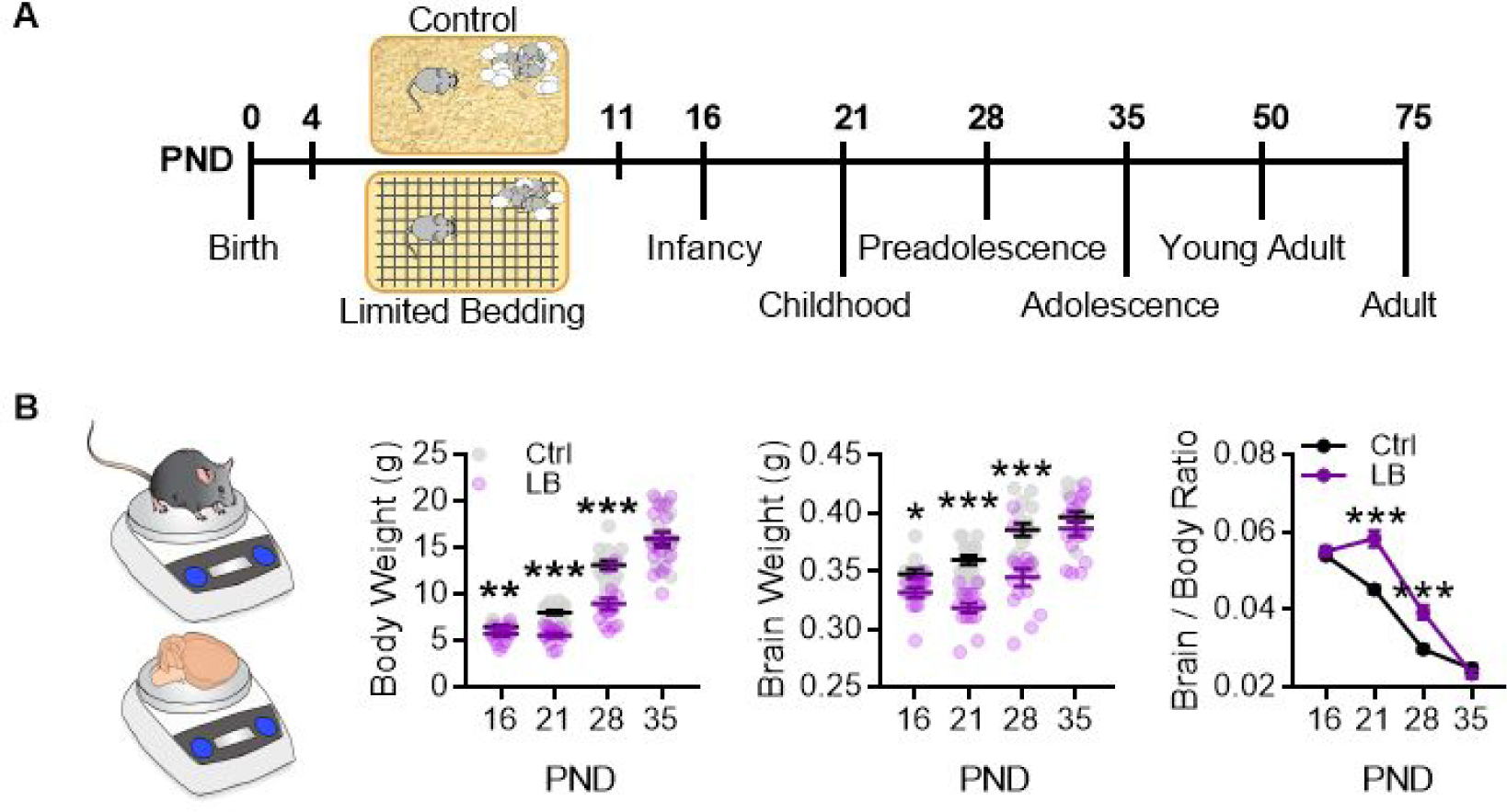
Limited bedding (LB) resources early in life, altered somatic and brain development. **A)** Timeline of resource restriction manipulation (PND 4-11) and description of developmental time points tested. **B)** Graph of whole body weight (left) brain weight (center) and the Brain to body weight ratio (right) as mice age from infancy (PND 16) into adolescence (PND 35). LB reared mice had a decrease in body weight at PND 16 (*p* = 0.0087), 21 (*p* < 0.001) and 28 (*p* < 0.001) with differences resolving by P35 (*p* = 0.86) (Ctrl n = 15, 18, 16, 16; LB n = 17, 16, 16, 16). Similarly, LB reared mice had decreased brain weight at PND 16 (*p* = 0.018), 21 (*p* < 0.001) and 28 (*p* < 0.001) with differences resolving by P35 (*p* = 0.22) (Ctrl n = 12,18,16,14; LB n = 14, 16, 16, 16). Mice reared in LB conditions had a higher brain to body weight ratio when compared to controls at PND 21 (*p* < 0.001) and 28 (*p* < 0.001). Dots in panel B represent individual data points. Bars represent group means +/− SEM. For B, Holm-Sidak multiple comparison analysis was used. * = *p* < 0.05, ** = *p* < 0.01, *** = *p* < 0.001.

### Deficits in fear recall during childhood

To determine if LB affected learning and expression of a conditioned fear memory, mice were conditioned to 6 tones (75db, 30 sec) each co-terminating with a 1 second foot-shock (0.57 mA). To dissociate cue-memory from context-memory, mice were habituated to two contexts (contexts A and B), then conditioned to a tone/shock association in context A and tested for memory recall in context B (**Figure 2A top**). LB reared mice showed significantly lower levels of freezing compared to control mice during cue recall at PND 22, but not at other developmental time points (**Figure 2A bottom**), indicating a possible delay in cue-associated fear learning in LB mice. To control for differences in conditioning, conditioning curves of control and LB mice were matched for levels freezing during acquisition (**Supplementary Figure 1 and Supplementary Figure 2**). The results showed that deficits in fear expression could not be solely explained by deficits in conditioning. To determine if the lower levels of freezing in LB mice at PND 22 was the result of deficits in acquisition, consolidation, or recall, separate cohorts of mice were conditioned at PND 21 and tested at either 1 hr (pre-consolidation), 6 hrs (post-consolidation), 24 hrs (short term recall), or 7 days (long term recall) post conditioning (**Figure 2B**). At 1 hr, there was a decrease in freezing observed in LB reared mice compared to controls. At 6 hrs, no differences in levels of freezing were detected between LB and control reared mice, indicating that LB mice could freeze to the cue and appeared to have consolidated the fear memory. By 24 hrs, the LB mice showed lower freezing compared to control mice, suggestive of deficits in fear memory. Interestingly, at 7 days post conditioning, LB freezing levels were back to control levels, suggesting that initial learning was intact. Together the results suggest that LB reared mice form a cue-associated fear memory at PND 21, but are not able to behaviorally express this memory during early development. A battery of tests were conducted to insure that effects of fear expression were not due to differences in anxiety-like behavior (light/dark box test, **Figure 2C**), locomotion (open field test, **Supplementary Figure 3**), or foot-shock sensitivity and reactivity (**Supplementary Figure 4**): In all of these tests, no differences were detected between LB and control at PND21.

**Figure 2.**
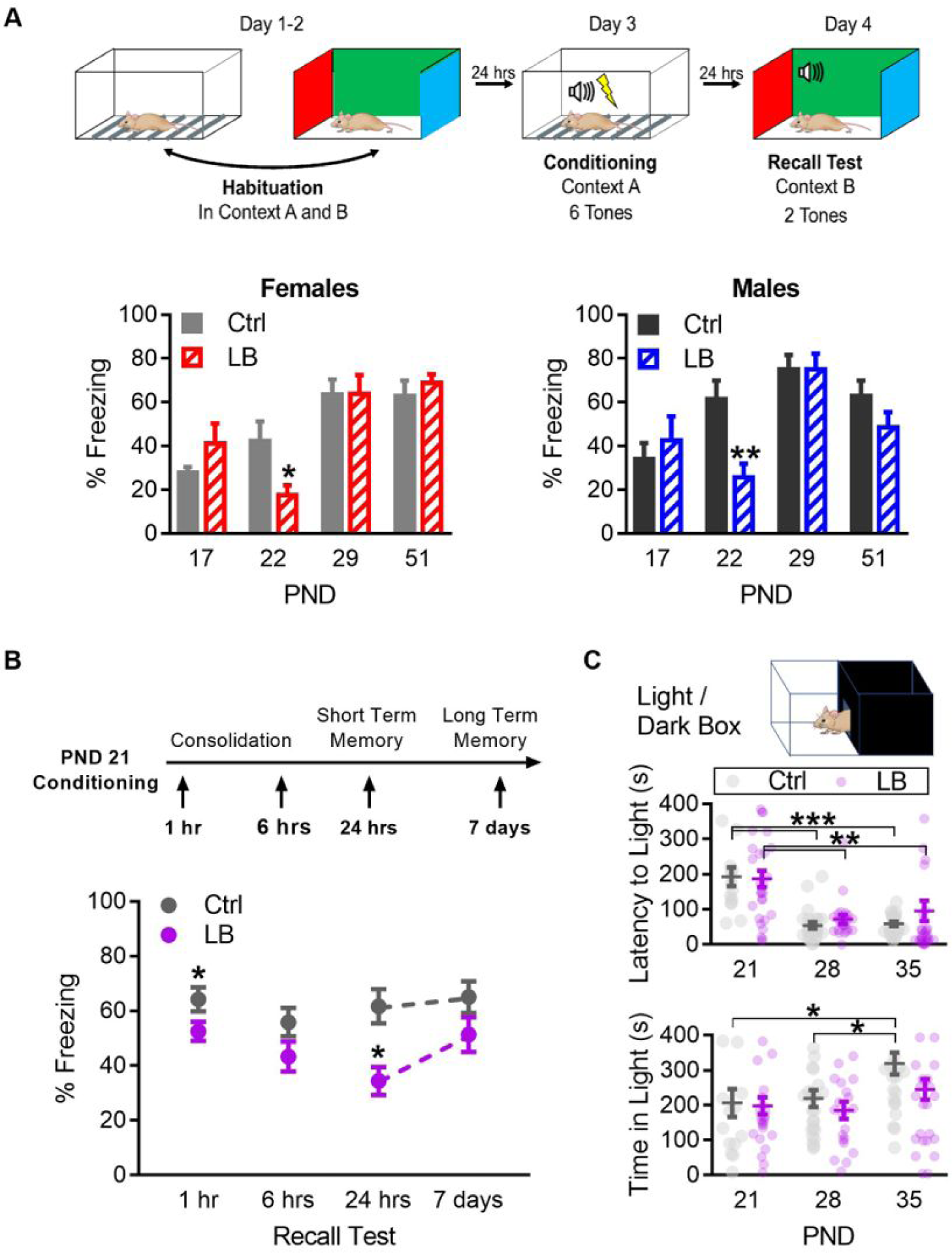
LB affects the short term expression of fear during early development. **A)** Schematic of auditory fear conditioning protocol (top), and graphs of the percent time female (bottom left) and male (bottom right) mice spent freezing (immobile) during the recall test. LB female (*p* = 0.041) and male (*p* = 0.0085) mice exhibited decreased fear expression at PND 22 when compared to age and sex matched controls, an effect not observed at other ages. (For females, Ctrl n = 11, 8, 12, 11; LB n = 11, 15, 9, 12. For males, Ctrl n = 16, 9, 12, 12; LB n = 10, 13, 11, 12). **B)** Schematic of experimental protocol (top). All mice were conditioned at PND 21 and tested at only one time point: 1 hr, 6 hrs, 7 hrs, or 7 days post conditioning. Graph depicting changes in freezing levels of distinct cohorts of mice during recall tests of varying delays (bottom). LB mice had decreased freezing at 1 hr (*p* = 0.038), 24 hrs (*p* = 0.002) but not 6 hrs (*p* = 0.10) or 7 days (*p* = 0.12) post conditioning. (Ctrl n = 34, 30, 18, 18; LB n = 42, 27, 17, 18). **C)** Depiction of the light/dark box used to assess anxiety-like behavior (top). Mice were placed in the dark side of the box. The latency to light (center) and the total time spent in the light side of the box (bottom) are shown. Total time spent in the light/dark box was 420 seconds. Age (*p* < 0.0001), but not rearing condition (*p* = 0.37), significantly affected the time it took mice to exit the dark side of the box (center). PND 21 mice of both LB and control reared conditions took more time to enter the light side of the box when compared to mice from the same rearing condition at PND 28 (LB *p* = 0.0005; Ctrl *p* = 0.0003) and PND 35 (LB *p* = 0.003; Ctrl *p* = 0.0005). Similarly, LB did not affect the time that mice spent in the light side of the light / dark box (*p* = 0.10; bottom) compared to controls. However, significant effects of age on the time spent in the light side of the box were observed (*p* = 0.0057). Specifically, control PND 35 mice spent significantly more time in the light side when compared to control reared mice aged PND 21 (*p* = 0.043) and PND 28 (*p* = 0.040). (Ctrl n = 15, 24, 23; LB n = 28, 22, 31). Bars represent group means +/− SEM. Dots in panel C represent individual data points. Unpaired two-tailed student t.tests were used in A and B. For C a two-way ANOVA followed by a Sidak’s multiple comparison analysis was used. * = *p* < 0.05, ** = *p* < 0.01, *** = *p* < 0.001.

### Accelerated Parvalbumin differentiation in the BLA, but not mPFC

To determine if the altered trajectory of fear expression may be due to alterations in the development of circuits supporting this behavior, we used various markers to track maturation of subclasses of neurons across distinct brain structures. We focused on the developmental trajectories of mPFC and BLA, structures known to be involved in emotional regulation and fear conditioning (Etkin et al., 2011; Sierra-Mercado et al., 2011; Arruda-Carvalho and Clem, 2015).

Using immunocytochemistry, we labeled Parvalbumin positive interneurons (PV+ cells), a late differentiating subclass of inhibitory interneurons (Rymar and Sadikot, 2007; Mukhopadhyay et al., 2009; Bartolini et al., 2013), in the brain of LB and control reared mice at PND 16, 21, 28, 50, and 75. Importantly, PV neurons are known to begin differentiating around PND 10-28, reaching maturation at approximately PND 30 (Berdel and Moryś, 2000; Dávila et al., 2008). In BLA, LB led to an increase in PV+ cell densities at PND 21 when compared to control reared mice at this age (**Figure 3A**). Importantly, no differences in PV+ cell densities were observed at PND 16, with both groups expressing low densities of PV+ cells. Furthermore, the difference in PV+ cell densities between LB and control subsided by PND 28, with control mouse PV+ levels rising to match those observed in LB mice (**Figure 3A left**). The increase in PV+ cell density at PND 21 was not observed in the other regions of the fear circuit assessed here, including the prelimbic (PL; **Figure 3A center**) or infralimbic (IL; **Figure 3A right**) subregions of the mPFC. These findings suggest that LB accelerates PV+ cell differentiation in the BLA but not the mPFC.

Next, Western blot analysis was used to determine if LB impacts protein levels for other classes of interneurons (calbindin/calretinin), glutamatergic neuronal markers (VGLUT1), or markers of myelination (myelin basic protein) across early developmental time points (**Figures 3B and 3C**). In BLA, LB had no effect on calbindin, VGLUT1, or myelin basic protein levels at any age tested, with only a modest but significant increase in calretinin levels at PND 16, that dissipated by PND 21 (**Figure 3B**). However, in mPFC, while LB showed an increase in calretinin and calbindin protein levels at PND 16 and increased myelin basic protein levels at PND 28, the only effect observed at P21 was a modest but significant reduction in calretinin protein levels (**Figure 3C**). Together our data shows that LB drives cell-type specific accelerated maturation of PV+ cells in BLA.

**Figure 3.**
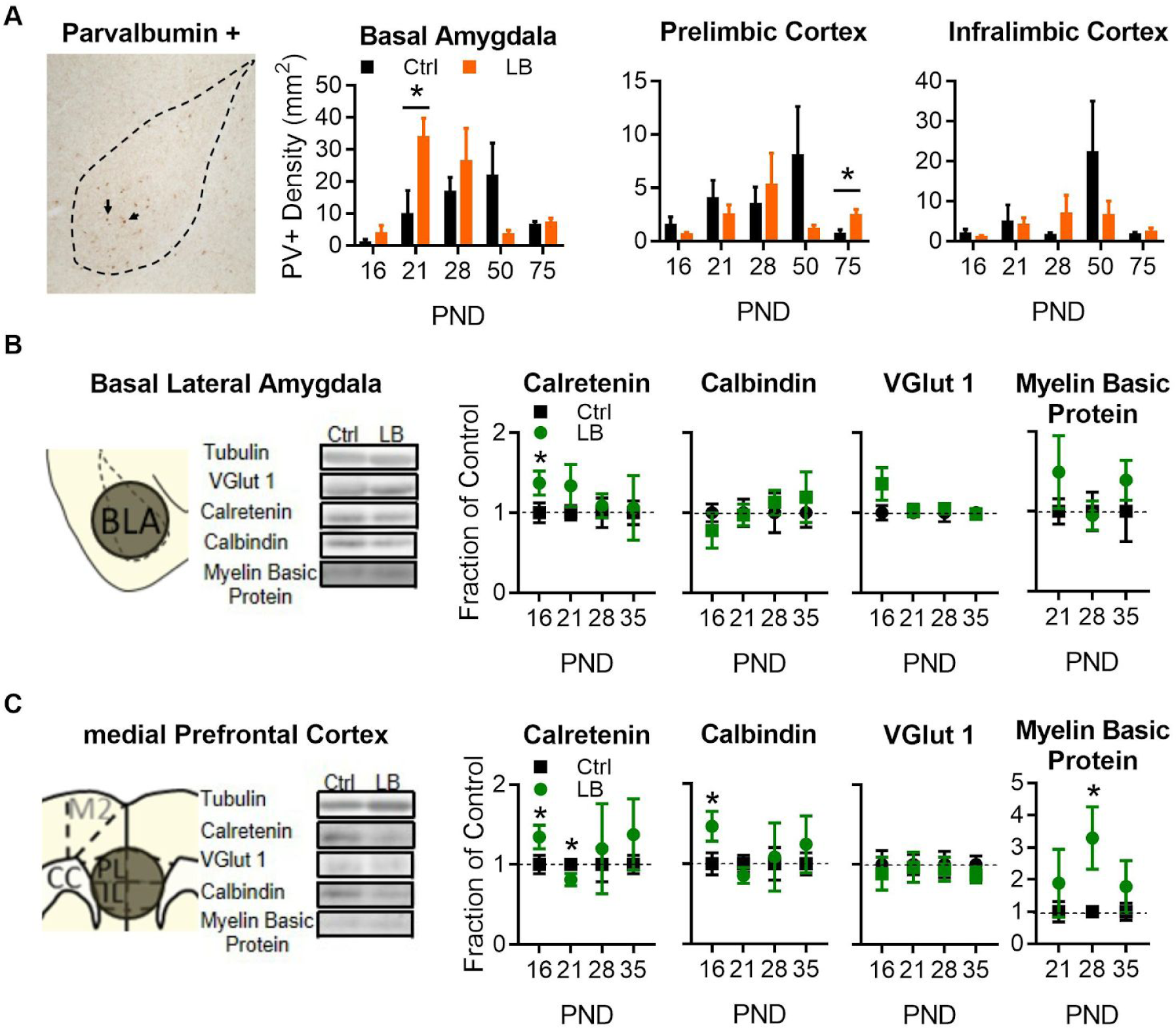
LB alters select neuronal populations in mPFC and BLA. **A)** Representative image of immunohistochemical stain in the BLA of an LB PND 21 mouse (left). From left to right, graphs showing the density of PV+ cells in the BLA, PL, and IL of mice at different ages. Immunohistochemical analysis revealed that LB significantly increased Parvalbumin + (PV+) neurons at PND 21 in the basolateral amygdala (*p* = 0.037) but not in the prelimbic cortex (*p* = 0.43) or in the infralimbic cortex (*p* = 0.87) at early developmental timepoints. Differences between LB and control mice did emerge in the prelimbic cortex at PND 75 (*p* = 0.019). For BLA (Ctrl n = 5, 7, 5, 6, 6; LB n = 5, 5, 5, 5, 5). For PL and IL (Ctrl n = 5, 6,6,7,6; LB n = 5, 6, 6, 6, 5). **B)** Western blot analysis of BLA tissue. Diagram and sample blots of PND 35 mice are shown. Graphs showing differing protein levels of mice from infancy into adolescence. LB increased calretinin levels (*p* = 0.021) at PND 16. No differences between LB and control mice were observed in BLA for calbindin, VGLUT1, or myelin basic protein levels. For calbindin, calretinin, and myelin basic protein (Ctrl n = 7, 7, 6, 7; LB n = 6, 6, 7, 7). For VGLUT1 (Ctrl n = 6, 7, 7, 7; LB n = 7, 7, 7, 7). **C)** Western blot analysis of mPFC tissue. Diagram and sample blots of PND 35 mice are shown. Graphs showing differing protein levels of mice from infancy into adolescence. LB increased calretinin (*p* = 0.041) and calbindin (*p* = 0.030) protein levels at PND 16. The PND 16 increase in calretinin protein levels was followed by a relative decrease in protein levels at PND 21 (*p* = 0.029). An increase in mPFC Myelin Basic Protein was observed exclusively at PND 28 (*p* = 0.028). For calbindin, calretinin, and myelin basic protein (Ctrl n = 6, 7, 6, 7; LB n = 6, 7, 5, 6). No effects of VGLUT1 were observed in the mPFC. For VGLUT1 (Ctrl n = 6, 7, 7, 7; LB n = 6, 7, 7, 7). Bars represent group means +/− SEM. Unpaired two-tailed t.tests between control and LB were used for A. For B and C, single sample two-tailed t.test analysis was used to determine if LB reared levels differed from the null hypothesis of no change (the value 1). * = *p* < 0.05, ** = *p* < 0.01, *** = *p* < 0.001.

### PV inhibition in BLA rescues fear expression deficits

To understand the neural basis of the suppressed fear expression at PND 21, we focused on the potential role of the premature differentiation of PV+ neurons in the BLA (**Figure 2A**). The focus on PV+ cells is related to previous reports that have demonstrated a role for BLA PV+ cells in modulating fear expression (Wolff et al., 2014; Davis et al., 2017). Therefore, we hypothesized that the early emergence of PV+ cells in the pre-adolescent BLA may be leading to the decreased fear expression observed at PND 22. To test this hypothesis, we used transgenic mice that selectively express the optogenetic construct halorhodopsin in PV+ cells. To generate mice, female mice homozygous for cre under the control of a parvalbumin promoter (PV-Cre +/+) were bred to male mice heterozygous for Floxed Halorhodopsin (NpHR +/−). The cross results in light control mice (PV-Cre (+/−) NpHR (-/-)) which do not express the halorhodopsin construct and PV Halo mice (PV-Cre (+/−) NpHR (+/−)) which express the halorhodopsin construct. Importantly mice that express halorhodopsin are also positive for EGFP, allowing us to verify ELA effects on cell density. This strategy allowed us to silence PV+ cells in a time and region specific manner.

To assess the replicability of ELA induced accelerated PV+ differentiation, the density of PV+ cells was quantified in control and LB reared mice from PV Halo mice (PV-Cre (+/−) NpHR (+/−)), which co-express an EGFP reporter. The increase in BLA PV+ in PND 21 LB mice replicated the prior immunohistochemical findings (**Figure 2A**). Importantly, at PND 21 LB PV Halo mice had a greater density of PV+ cells in the BLA compared to age matched control PV Halo mice (**Figure 4A**), effects not observed in PL or IL (**Figure 4B**).

**Figure 4.**
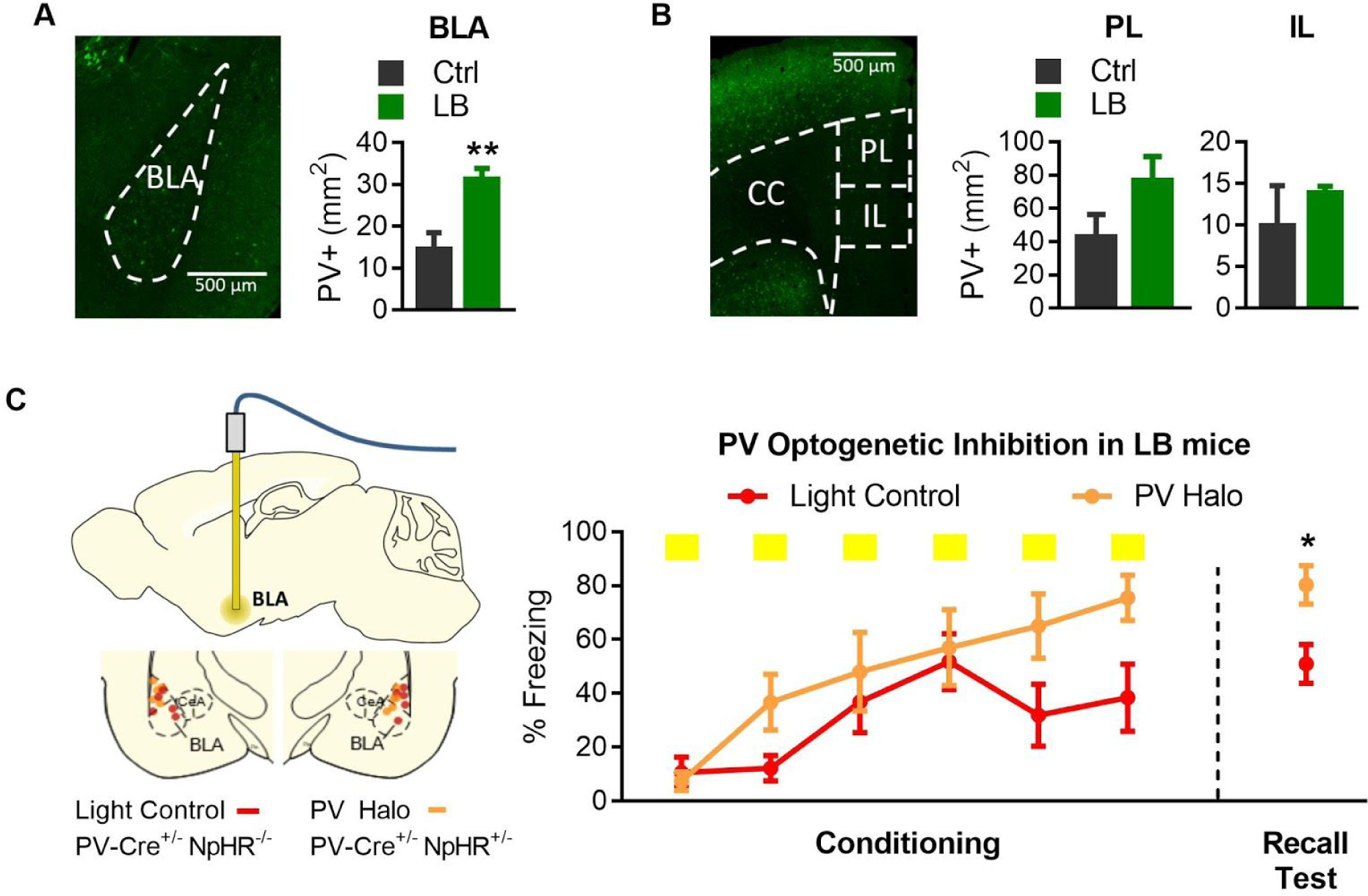
Optogenetic inactivation of parvalbumin (PV+) cells was sufficient to increase fear expression at PND 22 of LB mice. **A)** Representative image of PV+ cells in BLA of PV-Cre (+/−) NpHR (+/−) (“PV Halo”) mouse line (left). LB increased the density of PV+ cells in the BLA (*p* = 0.0081; n = 5 per group). **B)** Representative image of PV+ cells in the mPFC of a PV Halo (left). LB did not affect the density of PV+ cells in PL (*p* = 0.14) or IL (*p* = 0.45) compared with control reared mice (n = 4 per group). **C)** Graph showing conditioning and recall test. Optogenetic inhibition occurred during the conditioning tones. LB reared Light control (PV-Cre^+/−^ / Halo^−/-^) and PV Halo (PV-Cre^+/−^ / Halo^+/−^) mice were bilaterally implanted with an optical fiber into BLA, optic fiber placements are shown (left). Optogenetic inactivation of PV+ cells increased freezing during conditioning (*p* = 0.0064), and resulted in an increase in freezing on the recall test 24 hrs later (*p* = 0.018; For LB (light control) n = 5, for LB (PV Halo) n = 7. A two-way Anova was used to assess differences in fear conditioning, for all other analysis a two-tailed unpaired student t.tests were used. Bars represent group means +/− SEM. * = *p* < 0.05, ** = *p* < 0.01, *** = *p* < 0.001.

To test if inhibiting PV+ cells in the BLA of LB reared mice during fear learning could rescue the observed freezing deficits at PND 22, we optogenetically silenced PV + cells in the BLA during fear conditioning. Bilateral inactivation of PV+ cells within the BLA during the conditioning tones resulted in increased freezing 24 hrs later, during the recall test (**Figure 4C**). To ensure that the increased freezing was not due to effects on locomotion or anxiety-like behavior, mice were placed in an open field and PV+ cells in the BLA were optogenetically inhibited and the behavior of mice was tracked. No effects of optogenetic inhibition of PV+ cells were found for measures of anxiety-like behavior or general locomotor activity of mice (**Supplementary Figure 5A**). Thus, the increase in freezing behavior observed during recall was likely specific to the animals response to the conditioning paradigm. To insure that inhibition of PV+ cells was modulating BLA activity, a subset of unilaterally implanted mice were administered light for 15 minutes while freely moving within their homecage. Since parvalbumin neurons work to inhibit neuronal activity we hypothesized that inactivation of the inhibitory neurons should lead to increased excitation and therefore increase cFos. We found that mice exhibited increased cFos labeling in the optogenetically inhibited side when compared to the non-inhibited (naive) side (**Supplementary Figure 5B**). Further, electrophysiological control experiments have been carried out by our lab in the OFC of this same line of mice, and demonstrated robust inhibition PV+ cells in response to light (Goodwill et al., 2018). Together, these results suggest that the premature increase of PV+ cells in LB mice may be leading to increased inhibition of the BLA resulting in decreased freezing during the recall of conditioned fear during childhood.

**Figure 5.**
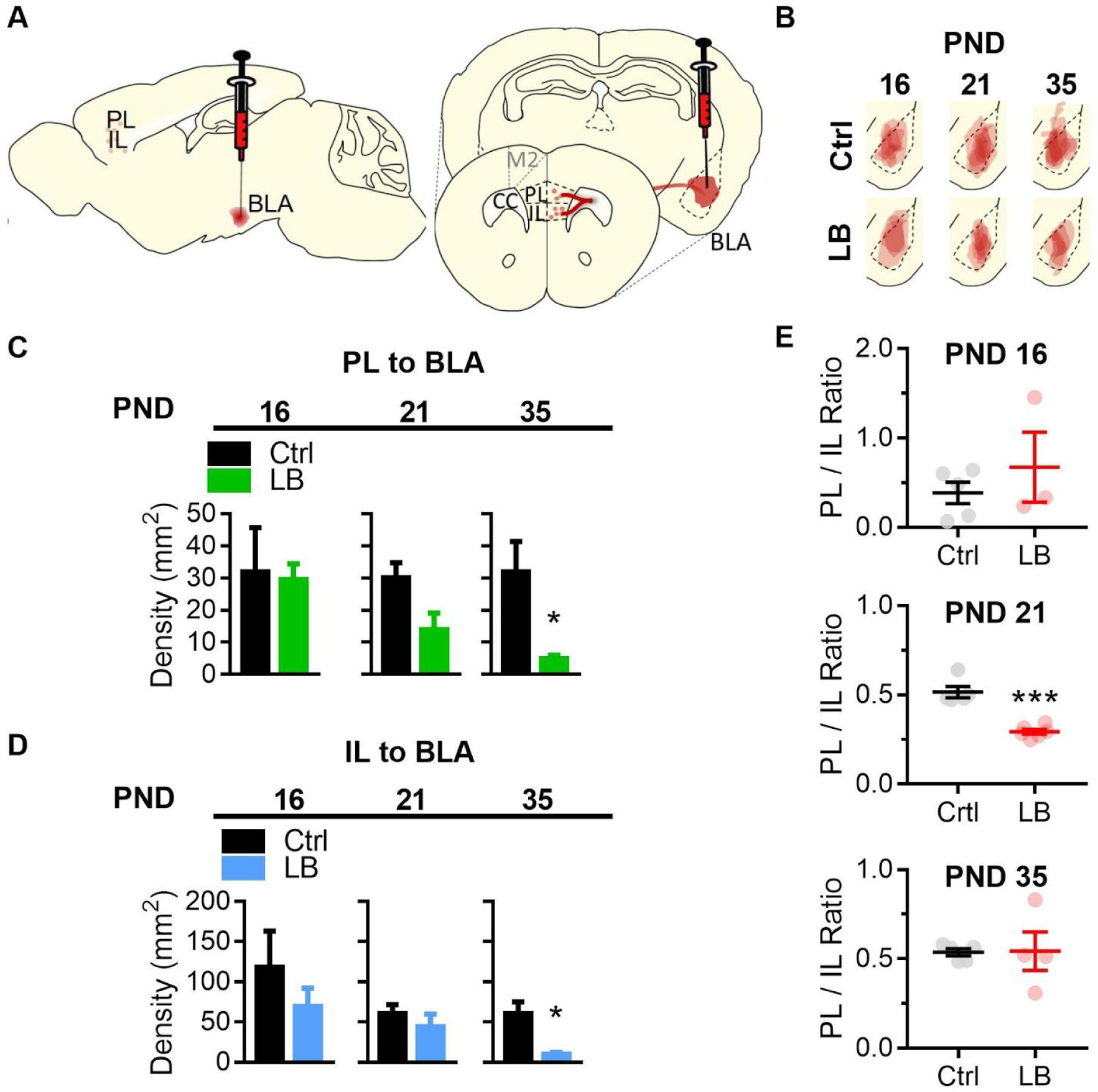
LB altered mPFC to BLA connectivity. **A)** Diagram of CTB 594 retrograde injection into the BLA. **B)** Tracings of CTB injections into the BLA. **C)** Graphs showing the differences in PL to BLA densities from infancy into adolescence. The density of PL to BLA projecting cells in PL did not differ at PND 16 (*p* = 0.90) or PND 21 (*p* = 0.052). However, LB significantly decreased the density of PL to BLA projections at PND 35 (*p* = 0.016) when compared to control mice. **D)** Graphs showing the differences in IL to BLA densities from infancy into adolescence. LB did not affect the density of IL to BLA projections at PND 16 (*p* = 0.45) or PND 21 (*p* = 0.42) but led to a significant decrease in LB relative to control at PND 35 (*p* = 0.023). **E)** Graphs showing the differences in PL/IL to BLA input ratios from infancy into adolescence. LB altered the balance of PL and IL projections to BLA. The ratio of the density of PL and IL to BLA cells was calculated. LB decreased the PL/IL ratio at PND 21 (*p* < 0.0001) but not at PND 16 (*p* = 0.41) or PND 35 (*p* = 0.94). Bars represent group means +/− SEM. Unpaired two-way student t.test were used to assess statistical significance between control and LB mice. For PND 16 (Ctrl n = 5, LB n = 3), PND 21 (Ctrl n = 5, LB n = 6), PND 35 (Ctrl n = 6, LB n = 4). * = *p* < 0.05, ** = *p* < 0.01, *** = *p* < 0.001.

### Resource insecurity decreased mPFC to BLA anatomical connectivity

We determined that LB affects mPFC and BLA cellular maturation in a marker and region specific manner. However, the impact of LB on mPFC and BLA connectivity remained unknown. To test if LB altered the timing and density of projections from mPFC to BLA, a retrograde tracer, cholera toxin B (CTB) was injected unilaterally into the BLA and the number of labeled cells in PL and IL were quantified. Injections were performed one day prior to the time point of interest (e.g. PND 15 for PND 16) and perfusions performed 1 day later (e.g. PND 17 for PND 16). Three time points were tested (PND 16, 21, and 35; **Figure 5A-B**). LB did not affect the density of PL to BLA or IL to BLA labeled cells at PND 16 or 21. However, LB reared mice did show a decrease in PL to BLA and IL to BLA projecting cells at PND 35 (**Figure 5C and 5D**). PL projections into BLA promote fear learning, while IL projections into BLA support fear extinction (Sierra-Mercado et al., 2011; Lee and Choi, 2012; Do-Monte et al., 2015a; Giustino and Maren, 2015). Next we sought to determine if LB altered the balance of PL and IL inputs into BLA across development. To assess PL/IL projection ratio for each mouse, the mean density of PL to BLA projecting cells was divided by the mean density of IL to BLA projecting cells (PL/IL; **Figure 5E**). LB reared mice had a significant decrease in PL/IL ratio compared to control reared mice, an effects only observed at PND 21. These results suggest that IL and PL projections to BLA are affected by LB and that LB may drive a mismatch in PL/IL balance over early development, with possible implications for expression of fear learning at PND 21.

## Discussion

Here, we provide a potential causal link between the neurodevelopmental and behavioral consequences of limited resource rearing on fear learning and developmental fear expression. We found that LB decreased somatic and brain growth during childhood and pre-adolescence, with more robust effects on the body than brain. Delayed brain growth was accompanied by significant effects on the maturational timing of subpopulations of cells in select brain regions as well as on regional connectivity. LB led to accelerated PV+ cell differentiation within the BLA, and altered the balance of mPFC inputs to BLA at PND 21 (childhood). Furthermore, at PND 21 LB reared mice were able to learn, but not express, an auditory conditioned fear response, which was rescued by inhibiting PV+ cells activity in the BLA of LB reared mice.

Previous research in humans has shown that childhood poverty, specifically during infancy, is associated with altered regional brain volume, with the prefrontal cortex and amygdala being affected (Sheridan et al., 2012; Hanson et al., 2013; Luby et al., 2013; Hair et al., 2015). However, the mechanisms by which poverty confers risk for altered brain development and how these factors impact the development of region and cell specific population of neurons remains largely unknown. Here, rearing mice under conditions of resource restriction, which may mirror aspects of poverty, altered the development of select populations of neurons in the mPFC and BLA, key regions involved in fear learning and threat assessment. In the BLA, LB accelerated PV+ neuronal differentiation and altered the timing, density, and PL/IL input balance to BLA. Further, LB altered calretinin and calbindin protein levels, as well as levels of myelin basic protein at select developmental time points in the mPFC, effects that may contribute to altered development and functional readouts for behaviors that were not assessed here. Consistent with our results, studies in humans have found that childhood poverty decreases prefrontal activity in adulthood and reduces the ability of prefrontal cortex to suppress amygdala activity (Kim et al., 2013).

Given the developmental changes observed in LB reared mice, we sought to test if these effects may be related to changes in threat assessment and responding to fear associated stimuli. As the mPFC and BLA are critical for fear learning, mice at different developmental ages underwent Pavlovian tone-shock association learning (Anglada-Figueroa and Quirk, 2005; Etkin et al., 2011; Sierra-Mercado et al., 2011; Arruda-Carvalho and Clem, 2015; Giustino and Maren, 2015). Control reared mice showed the expected developmental trend of being able to condition to a tone-shock association at PND 21, but not at PND 16. Interestingly, LB reared animals were unable to express the conditioned memory 24 hr after conditioning at either PND 16 or PND 21, indicating a later developmental emergence of this behavior. Importantly, LB mice were able to express the conditioned memory at a short delay (6 hrs), indicating that deficits in freezing were not due to an inability to engage the freezing response. Furthermore, the freezing response re-emerged 7 days following conditioning, indicating that learning had occurred. Thus, LB may disrupt the developmental ability of animals to express, but not form, specific types of conditioned fear memories.

Previous research from our lab has demonstrated that LB can induce accelerated PV+ differentiation in the hippocampus (Bath et al., 2016). Consistent with this prior work, we found that LB led to an earlier rise in PV+ cell density in BLA, with PV+ cell density being higher in LB reared mice compared to controls at PND 21. PV+ neuronal activity in the BLA has been shown to be capable of modulating fear expression (Wolff et al., 2014; Davis et al., 2017). In addition, PV+ cell density has previously been shown to negatively correlate with innate threat responses in post-weaned rats (Santiago et al., 2018). Consistent with the previously described role of PV+ neurons in BLA, we found that mice reared under LB conditions had decreased fear responding, as indexed by freezing to a tone previously paired with a shock at PND 21. We posit that the precocious maturation of PV+ cells in BLA may be blunting the ability to express, but not learn a fear association. In support of this interpretation of the data, optogenetic inhibition of PV+ cells in BLA was sufficient to rescue the low freezing phenotype of PND 21 LB reared mice. This work significantly extends our knowledge of the role of PV+ cells in fear expression, and possible mechanisms through which ELA can drive developmental changes in threat assessment and fear expression.

We show that LB increased PV+ cell differentiation, and that this effect may be driving the decrease in fear expression at PND 21. However, it is also possible that LB induced changes in mPFC to BLA connectivity may be contributing to the decreased freezing phenotype observed at PND 22. Previous studies in rats have shown that the mPFC can modulate auditory fear conditioning as early as PND 24, but not at PND 17 (Kim et al., 2009). Furthermore, it has been shown that stimulation of the IL subregion of the mPFC can decrease fear expression (Do-Monte et al., 2015a). Here, we found that at PND 21 there is a decrease in the ratio of PL/IL inputs to BLA, possibly resulting in more robust IL control of the BLA. Additional experiments will be required to determine if LB effects on IL projections into BLA or PL/IL balance are also contributing to altered processing of threat signals at this age.

In sum, this work reveals complex effects of limited resource rearing, a poverty-like model, on the timing of neuronal and circuit maturation and the developmental expression of fear learning. We found that resource restriction during early development can drive regional and cell selective effects on maturation, with profound implications for behavioral development. Specifically, we have found that LB accelerated the differentiation of PV+ neurons in BLA, and transiently altered the ability of mice to express a fearful memory. This transient inability to access and drive behavioral responding to fear-associated stimuli could have implications for risk taking behavior or behavioral manifestation of symptoms associated with early life traumatic events. The transient suppression of threat-associated fear behavior may increase risk taking (Haushofer and Fehr, 2014), increasing the chances of incurring a secondary stressor. Further, such observations may explain the latent period of neuropsychiatric symptom expression in children exposed to early life trauma (Teicher et al., 2009). Future studies assessing the contribution of altered development of these fear circuits to later symptom development will be needed.

## Author Contributions

GMN designed experiments in consultation with KGB. GMN carried out all behavioral, molecular, and immunohistochemical experiments. GMN and MB conducted CTB tracing experiments. GMN wrote the manuscript with KGB, MB reviewed and edited.

## Acknowledgements

This work was supported by funding by the National Institutes of Health, from the NIMH (MH115914-KGB; MH115049-KGB) and NINDS (NS105219-GMN). The authors would like to thank the Connors lab and the Moore lab at Brown University for facilitating access to the transgenic mice. We would like to thank Dr. Rebecca Burwell for permitting us the use of her lab’s microscope. In addition, the authors would like to thank Saba Nur Baskoylu for teaching GMN how to perform Western blots and assisting with collection and analysis of the shock sensitivity assay.

## Subjects

Approximately 1000 C57BL/6N wildtype and 45 transgenic, male and female, mice were used in this study. Original breeding stock was ordered from Charles River Labs. All wild-type C57BL/6N mice were bred in house. For optogenetic experiments, “PV-Cre” (JAX#008069) and “floxed Halo” (JAX#014539) mouse lines were derived from a breeding stock acquired from Jackson laboratories. All animals were housed according to NIH guidelines and maintained on a 12 hour light:dark cycle. Mice had free access to food and water throughout the study. All animal procedures were approved by the Brown University Institutional Animal Care and Use Committee and consistent with the National Institutes of Health Guide for the Care and Use of Laboratory Animals.

## Fragmented Maternal Care

Dam and pups were placed in low bedding conditions with limited access to nesting material for 7 consecutive days (PND 4 through PND 11) as previously described (Rice et al., 2008; Bath et al., 2016; Manzano-Nieves et al., 2018). Standard reared mice (designated as Controls or Ctrl) were left undisturbed in a standard home cage until weaning. Mice tested at PND 21 were weaned at PND 22.

## Fear Conditioning

During fear conditioning mice were presented with 6 tones (30 seconds, 4 KHz, 75 dB) with each tone co-terminating with a 1 second foot-shock (0.57 mA). Tone shock pairings were separated by an inter-trial interval of 1.5 minutes. Conditioning chamber specifications have previously been described (Manzano-Nieves et al., 2018).

## Light / Dark Box

Mice were tested in a Light / Dark Box that was built in house. To begin a trial, mice were placed in the dark side of the box which was connected to a light side by a small opening (~6 x 6 cm). To increase the brightness of the light side, a lamp, pointing toward the light side, was mounted on the lid of the dark side. The dimensions of the dark and light side of the chamber were the same and measured (height = 23 cm, width = 22 cm, length = 26 cm). A trial lasted a total of 10 minutes. Activity of the mouse during a trial was recorded and analyzed using Ethovision XT 11.0 software, with latency to first exit being hand scored by an observer blind to sex, age, and condition.

## Open Field Test

To test for differences in locomotor activity and anxiety-like behavior, mice were placed in an open field arena as previously described (Goodwill et al., 2019). Distance moved and the time spent in center of the arena were recorded during a 7 (**Extended Figure 3**) or 5 (**Figure 5**) minute test using the Ethovision video-tracking system.

## Shock Sensitivity Assay

Mice were exposed to a series of foot shocks, beginning at 0.06 mA and increasing at 0.02 mA intervals as previously described (Manzano-Nieves et al., 2018).

## Immunohistochemistry

For PV+ cell labeling, a rabbit anti-parvalbumin antibody (1:1,000; Millipore) was used. For c-Fos labeling a rabbit anti c-Fos (1:20,000; Millipore) was used to label brains as previously described (Bath et al., 2016).

## Western Blot

Standard protocol was followed. Primary antibodies used: mouse anti-Tubulin (1:2000, Cell Signaling Technology), rabbit anti-GAPDH (1:2000, Cell Signaling Technology), mouse anti-Calretinin (1:500, Swant), mouse anti-Calbindin (1:500, Swant) rabbit anti-Myelin Basic Protein (1:1000, Abcam), rabbit anti-VGLUT1 (1:1000, Millipore). Secondary antibodies used for this study were: HRP conjugated donkey anti-mouse (1:2000, Jackson immunoresearch) and donkey anti-rabbit (1:2000, Jackson Immunoresearch).

## Cholera toxin B Injections

Alexa 594 conjugated cholera toxin B (CTB) (Fisher Scientific) was used to retrogradely label the projections from PL and IL to BLA, in male mice, at postnatal ages PND 16, 21, and 35. CTB (1.0 mg/mL) was injected (0.15 ul) into the left BLA. Coordinates used for the CTB injections were as follows: (PND 15: DV = −5.075, ML = −3.05, AP = −1.1; PND 20: DV = −5.1, ML = −3.1, AP = −1.15; PND 34: DV = −5.2, ML = −3.15, AP = −1.2). Mice were perfused 48 hrs post injection.

## Optogenetic Inhibition of PV+ cells

Mice were bilaterally implanted on PND 15 (Placements: DV = −5.1, ML = = +/− 3.1, AP = −1.15), and began fear conditioning protocol at PND 19 (as described above), with optogenetic inhibition occurring at PND 21 (**Figure 5**). Photo-inhibition consisted of constant light (620 nm Plexbright LED, Plexon, Dallas, TX), driven by an LED driver (Plexon, Dallas, TX) during the 30 seconds of the tone (including the 1 sec foot-shock). Light power delivered, as measured through the optic fiber pre-implant, ranged from 1.5 – 2 mW per side.

## Microscopy Analyses

Neurolucida software was used to analyze immunohistochemical data. Either a light (for DAB staining) or epi-fluorescent microscope (for fluorescence) was used when appropriate. For each brain region 3-4 sections per brain were averaged to obtain a mean density.

## Statistical Analyses

Statistical analysis used is reported in figure legends. For CTB analysis the predicted values from an ANCOVA were used to correct for the placement and size of the injection. The area of the BLA and the area of the injection at the site of the injection were used as covariates. SPSS statistical analysis software was used to acquire the predicted values. Statistical analysis was performed using Prism Graphpad statistical and graphing software.

## Supplementary Methods

### Subjects

Approximately 1000 C57BL/6N wildtype and 45 transgenic, male and female, mice were used in this study. Original breeding stock was ordered from Charles River Labs. All wild-type C57BL/6N mice were bred in house. For optogenetic experiments, “PV-Cre” (JAX#008069) and “floxed Halo” (JAX#014539) mouse lines were derived from a breeding stock acquired from Jackson laboratories. For optogenetic experiments homozygous PV-Cre mice were bred with heterozygous floxed Halo mice resulting in 2 groups of offspring, PV-Cre^+/−^ / Halo^−/-^ (Light Controls) and PV-Cre^+/−^ / Halo^+/−^ (PV Halo). All animals were housed according to NIH guidelines and maintained on a 12 hour light:dark cycle. Mice had free access to food and water throughout the study. All animal procedures were approved by the Brown University Institutional Animal Care and Use Committee and consistent with the National Institutes of Health Guide for the Care and Use of Laboratory Animals.

### Fragmented Maternal Care

LB was modeled through a resource restriction paradigm, in which dam and pups were placed in low bedding conditions with limited access to nesting material for 7 consecutive days (PND 4 through PND 11). This manipulation results in a fragmentation in maternal care (Rice et al., 2008; Bath et al., 2016). Four days after the birth of a litter (PND 4), the dam and pups were transferred from their standard home cage with cob bedding and a 4 x 4 cm cotton nestlet to an LB cage containing a wire mesh floor and a 2 x 4 cm cotton nestlet. The mice continued to have *ad libitum* access to food and water. Following one week (PND 11), pups and dams were returned to their standard housing with full bedding and nesting material. Standard reared mice (designated as Controls or Ctrl) were left undisturbed in a standard home cage until weaning. All pups were weaned and sex segregated at PND 21, with one exception. Mice tested at PND 21 were weaned following the completion of fear conditioning experiments at PND 22.

### Fear Conditioning

Fear conditioning was carried out in Med Associates (St. Albans City, VT) operant chambers. On days 1 and 2 mice were habituated to two distinct (differing in color, texture, and smell) chambers. Habituation trials lasted 5 minutes per chamber and were counterbalanced. On day 3, mice received tone-shock associative learning in the fear conditioning chamber. During fear conditioning mice were presented with 6 tones (30 seconds, 4 KHz, 75 dB) with each tone co-terminating with a 1 second foot-shock (0.57 mA). Tone shock pairings were separated by an inter-trial interval of 1.5 minutes. Testing for fear expression occurred on day 4 of the testing protocol, unless otherwise stated. Fear expression testing consisted of exposing mice to 2 tones in the control chamber (habituated chamber where no fear conditioning occurred). Different cohorts of animals, across multiple litters were used to test mice at the different developmental time points. Freezing behavior was scored automatically by the activity tracker module in Noldus Ethovision XT 11.0 and verified from video by observers blind to treatment and condition but not to age (as age could be inferred based upon differences in mouse size).

### Open Field Test

To test for differences in locomotor activity and anxiety-like behavior, mice were placed in an open field arena as previously described (Goodwill et al., 2019). Distance moved and the time spent in the center of the arena were recorded during a 7 (**Extended Figure 3**) or 5 (**Figure 5**) minute test using the Ethovision video-tracking system. The arena was digitally divided into 2 zones (center and periphery), as previously described (Goodwill et al., 2019). Decreased time in the center was used as a indicator of anxiety-like behavior.

### Shock Sensitivity Assay

To assess the minimum foot-shock intensity required to elicit a behavioral response (visible flinch or audible vocalization), PND 21 mice were placed in an operant conditioning chamber (Med associates, Fairfax, VT, USA). Mice were exposed to a series of foot shocks, beginning at 0.06 mA and increasing at 0.02 mA intervals. Each shock intensity was presented 3 times. The amplitude of the foot-shock at which a given mouse first flinched, and/or audibly vocalized to 2 out of 3 foot shocks at a given intensity was recorded by two independent observers blind to condition to insure agreement on these measures. Flinching was defined as the mouse moving it’s body reflexively downward, making its body smaller, directly following a foot-shock. Vocalization was defined as the emittance of an audible sound.

### Immunohistochemistry

To assess the relative density of PV+ cells and c-Fos expressing neurons, immunohistochemistry was performed on control and LB mice on the days stated in each experiment. Briefly, mice were deeply anesthetized with pentobarbital, transcardially perfused with buffered saline followed by 4% paraformaldehyde, and processed for immunohistochemistry as previously described (Bath et al., 2016). For PV+ cell labeling, a rabbit anti-parvalbumin antibody (1:1,000; Millipore) was used. For c-Fos labeling a rabbit anti c-Fos (1:20,000; Millipore) was used. Brain sections (40 µm) were mounted on charged glass slides, counter stained using a Hema 3 staining set (Fisher Scientific Company), dehydrated, and coverslipped for imaging.

### Western Blot

Male mice were sacrificed, brains were quickly dissected, weighed, and flash frozen on dry ice. The medial prefrontal cortex (mPFC), and the basolateral amygdala (BLA) were dissected and stored at −80ºC until processing.

Tissues were homogenized in RIPA buffer (with 1% protease and phosphatase inhibitor cocktail, Fisher Scientific) and supernatant was collected following centrifuging at 14,000 rpm at 4ºC for 10 minutes. Protein concentration was determined with a bicinchoninic acid (BCA) kit (Thermo Scientific, Waltham, MA). Protein lysates were each diluted to 1.0 mg/mL with RIPA buffer, heated at 90ºC for 10 minutes, and proteins separated by gel electrophoresis on a 12% sodium dodecyl sulfate–polyacrylamide gel electrophoresis (SDS-PAGE) gel, and transferred to polyvinylidene difluoride (PVDF) membrane. For the remaining portion of the Western blot protocol, transfer membranes were kept at 4ºC.

Membranes were blocked for 1 hour in 5% non-fat milk in Tris-buffered saline Tween-20 (TBST, containing 10 mM Tris, 150 mM NaCl, and 0.1% Tween-20, pH 7.6), followed by incubation with primary antibodies diluted in 5% non-fat milk / 0.5% bovine serum albumin in TBST at 4ºC overnight. Membranes were washed with TBST three times (15 minutes per wash), and incubated with secondary antibody in 5% non-fat milk / 0.5% bovine serum albumin in TBST for 1 hour. Membranes were then washed with TBST three times (15 minutes per wash), then visualized with Amersham ECL Western Blotting Detection Reagent (RPN2106, GE Life Sciences) using a C600 Azure Biosystems imaging system (Dublin, CA). Densitometry analysis was conducted with gel imaging module of NIH ImageJ software.

Primary antibodies used: mouse anti-Tubulin (1:2000, Cell Signaling Technology), rabbit anti-GAPDH (1:2000, Cell Signaling Technology), mouse anti-Calretinin (1:500, Swant), mouse anti-Calbindin (1:500, Swant) rabbit anti-Myelin Basic Protein (1:1000, Abcam), rabbit anti-VGLUT1 (1:1000, Millipore). Secondary antibodies used for this study were: HRP conjugated donkey anti-mouse (1:2000, Jackson immunoresearch) and donkey anti-rabbit (1:2000, Jackson Immunoresearch).

### Cholera toxin B Injections

Alexa 594 conjugated cholera toxin B (CTB) (Fisher Scientific) was used to retrogradely label the projections from PL and IL to BLA, in male mice, at postnatal ages PND 16, 21, and 35. CTB (1.0 mg/mL) was injected (0.15 ul) into the left BLA one day prior to the time-point of interest (e.g. PND 15 for PND 16). In order to inject the CTB into BLA across development separate coordinates were used for each age. Developmentally appropriate coordinates were empirically derived from pilot surgeries. Coordinates used for the CTB injections were as follows: (PND 15: DV = −5.075, ML = −3.05, AP = −1.1; PND 20: DV = −5.1, ML = −3.1, AP = −1.15; PND 34: DV = −5.2, ML = −3.15, AP = −1.2). Mice were perfused 48 hrs post injection. Brain were dissected, sectioned (40 µm), mounted, counter-stained with DAPI (Fisher Scientific) and visualized using a fluorescent microscope. The density of CTB positive cells in the PL and IL was measured.

### Optogenetic Inhibition of PV+ cells

Female mice homozygous for Cre under the control of a parvalbumin driver (PV-Cre, JAX#008069) were crossed with a male heterozygous for the eNpHR3.0-EYFP fusion protein (NpHR)-EYFP (JAX#014539) with an upstream floxed stop cassette. The cross resulted in the selective expression of halorhodopsin in PV+ cells (PV-Cre ^+/−^, Halo ^+/−^ “PV Halo mice”) and mice from the same litter that were Cre positive, but lacked the optogenetic channel(PV-Cre^+/−^, Halo ^−/-^ “Light control”). Mice were bilaterally implanted on PND 15 (Placements: DV = −5.1, ML = = +/− 3.1, AP = −1.15), and began fear conditioning protocol at PND 19 (as described above), with optogenetic inhibition occurring at PND 21 (**Figure 5**). During conditioning, PV+ cells in the BLA were photo-inhibited with constant light (620 nm Plexbright LED, Plexon, Dallas, TX), using an LED driver (Plexon, Dallas, TX) during the 30 seconds of the tone (including the 1 sec foot-shock). The light power delivered, as measured through the optic fiber pre-implant, ranged from 1.5 – 2 mW per side. Following the fear conditioning protocol, a subset of the mice were tested for locomotion in an open field under conditions of light stimulation.

### Microscopy Analyses

Neurolucida software was used to analyze immunohistochemical data. Either a light (for DAB staining) or epi-fluorescent microscope (for fluorescence) was used when appropriate. For quantification of neuronal cell density, brain regions were traced at 4x magnification and borders were defined as shown in Paxinos and Franklin mouse brain atlas. Immune reactive or fluorescent positive neurons within each region were identified by an observer blind to condition and treatment (10x). All region contours with identified cells were saved and the number of cells and area within each contour was assessed using StereoInvestigator. For each brain region 3-4 sections per brain were averaged to obtain a mean density.

### Statistical Analyses

A two-tailed student’s *t-test* was used to compare between two groups. When more than two groups were assessed the appropriate ANOVA was performed. All ANOVA tests were followed by Sidak’s multiple comparison test, assessing the effects of treatment at each given age and/or a Tukey’s post-hoc comparison to assess developmental differences within each treatment. For all experiments, sex differences were tested for using the appropriate ANOVA analysis or t-test. For CTB analysis the predicted values from an ANCOVA were used to correct for the placement and size of the injection. The area of the BLA and the area of the injection at the site of the injection were used as covariates. SPSS statistical analysis software was used to acquire the predicted values. All data is presented in graphs and figures as mean ± SEM Statistical analysis was performed using a Prism Graphpad statistical and graphing software, statistical significance was described as *p* < 0.05.

