## Supplementary Figure 1 for "Early life adversity decreases fear expression in pre-adolescence by accelerating amygdalar parvalbumin cell development"

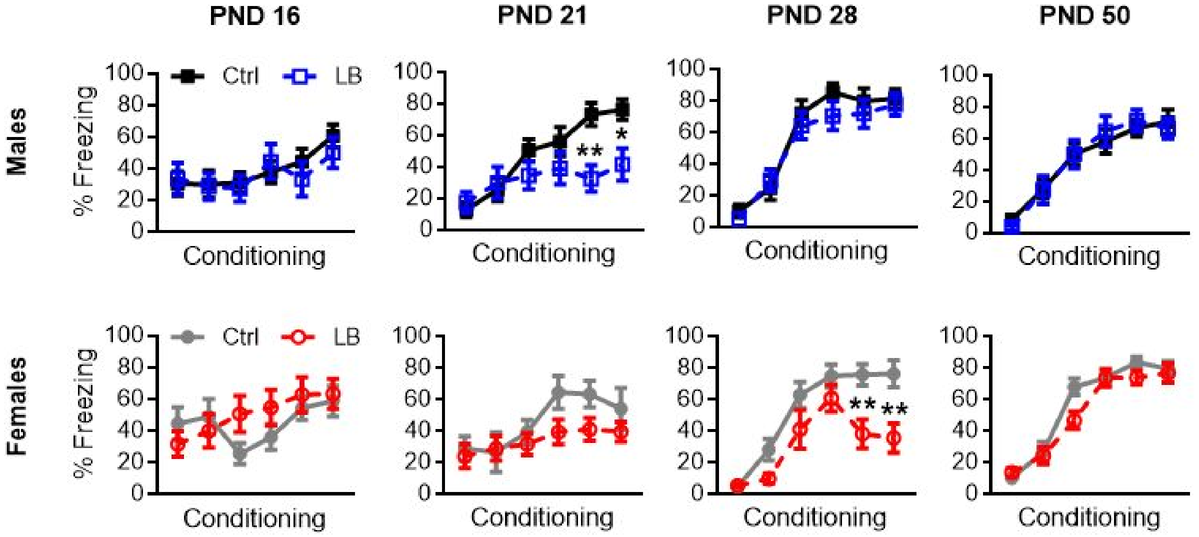


**Supplementary Figure 1.** LB affects conditioning in a sex and time specific manner. Graphs showing conditioning curves for males and females across development. Conditioning curves correspond to recall graphs in Figure 2. LB decreased conditioning to a tone/shock association in both males ($p$ = 0.0015) and females ($p$ = 0.020). Males did not show altered conditioning at PND 16, 28 or 50 because of rearing. However, LB decreased conditioning in females at PND 28 ($p$ < 0.0001), but not at 16 or 50. For females (Ctrl n = 11, 8, 12, 11; LB n = 11, 15, 9, 12. For males (Ctrl n = 16, 9, 12, 12; LB n = 10, 13, 11, 12). For all graphs the mean and SEM are presented. A two-way ANOVA followed by Sidak's multiple comparison analysis was used to assess differences in fear conditioning. A two-tailed paired student t.test was used to assess differences in recall test. * = $p$ < 0.05, ** = $p$ < 0.01, *** = $p$ < 0.001.
