## Supplementary Figure 2 for "Early life adversity decreases fear expression in pre-adolescence by accelerating amygdalar parvalbumin cell development"

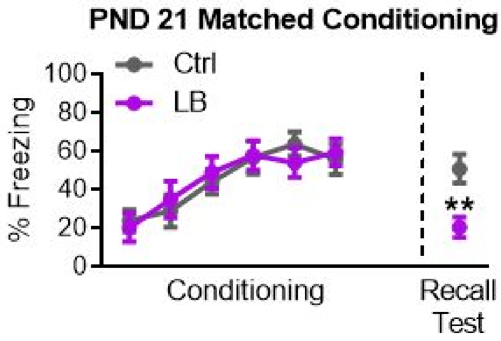


**Supplementary Figure 2.** Effect on recall at PND 21 are not a consequence of conditioning deficits. Graph of freezing matched for conditioning. Matching for conditioning did not alter effects on recall ($p$ = 0.0025). For Ctrl n = 13, for LB n = 14. A two-way ANOVA followed by Sidak's multiple comparison analysis was used to test for differences in fear conditioning. A two-tailed paired student t.test was used to test for differences in the recall test. * = $p < 0.05$, ** = $p < 0.01$, *** = $p < 0.001$.
