## Supplementary Figure 3 for "Early life adversity decreases fear expression in pre-adolescence by accelerating amygdalar parvalbumin cell development"

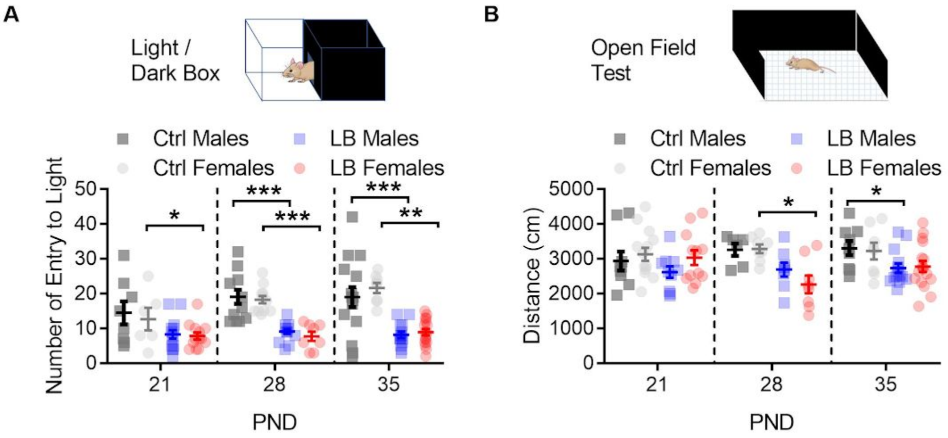


**Supplementary Figure 3.** LB did not affect anxiety like behavior or locomotion at PND 21. **A)** Female LB reared mice had a decrease in the number of entries into the light side of the box at PND 21 ($p$ = 0.046), 28 ($p$ < 0.0001), and 35 ($p$ = 0.0016). LB reared males had a decrease in the number of entries to the light side of the box at PND 28 ($p$ < 0.0001) and PND 35 ($p$ < 0.0001). (Ctrl males n = 8, LB males n = 14, Ctrl females n = 6, LB females n = 13), PND 28 (Ctrl males n = 12, LB males n = 14, Ctrl females n = 12, LB females n = 8), PND 35 (Ctrl males n = 15, LB males n = 14, Ctrl females n = 8, LB females n = 17). **B)** LB did not affect locomotion at PND 21, but did decrease locomotion at PND 28 in females ($p$ = 0.0029) and at PND 35 for males ($p$ = 0.022). PND 21 (Ctrl males n = 9, LB males n = 10, Ctrl females n = 14, LB females n = 12), PND 28 (Ctrl males n = 6, LB males n = 8, Ctrl females n = 8, LB females n = 8), PND 35 (Ctrl males n = 9, LB males n = 14, Ctrl females n = 8, LB females n = 17). For all graphs the mean and SEM are presented. A two-tailed paired student t.test was used to test for significance. * = $p$ < 0.05, ** = $p$ < 0.01, *** = $p$ < 0.001.
