## Supplementary Figure 4 for "Early life adversity decreases fear expression in pre-adolescence by accelerating amygdalar parvalbumin cell development"

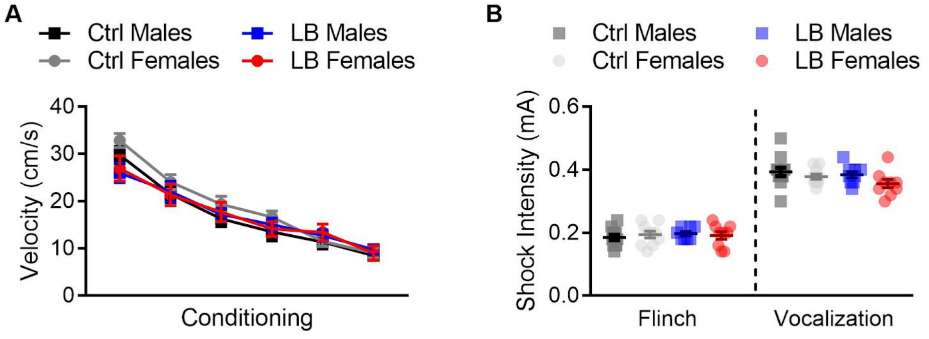


**Supplementary Figure 4.** LB did not affect somatosensation at PND 21. **A)** To assess if LB affected the response of mice to the conditioning foot-shocks, we plotted the mean velocity of mice during the one second foot-shock. We found that LB did not affect the mean velocity at which mice responded to the foot-shocks. (Ctrl males n = 39, LB males n = 38, Ctrl females n = 39, LB females n = 30). **B)** At PND 21, no effects of LB were observed on the minimum foot-shock intensity required for mice to flinch or audibly vocalize in response to the foot-shocks. (Ctrl males n = 12, LB males n = 9, Ctrl females n = 10, LB females n = 9). For all graphs the mean and SEM are presented. A two-way ANOVA followed by Sidak's multiple comparison analysis was used to assess differences in shock reactivity during conditioning. A two-tailed paired student t.test was used to assess differences in flinching and vocalization. * = $p < 0.05$, ** = $p < 0.01$, *** = $p < 0.001$.
