## Supplementary Figure 5 for "Early life adversity decreases fear expression in pre-adolescence by accelerating amygdalar parvalbumin cell development"

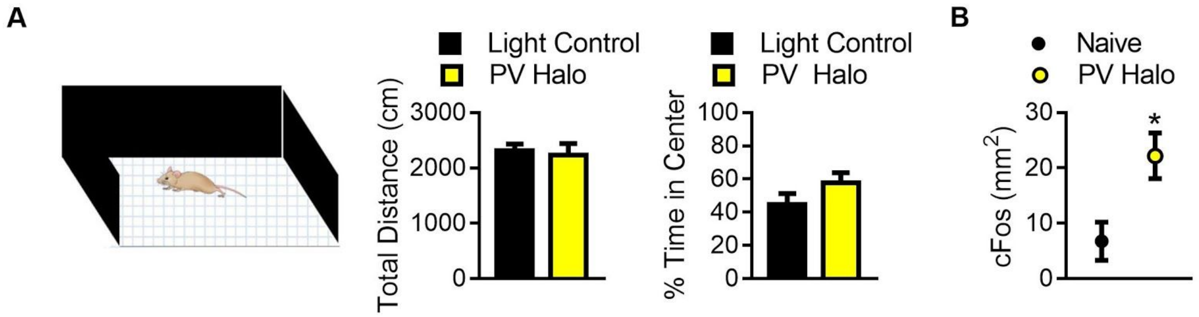


**Supplementary Figure 5.** Optogenetic inhibition did not affect locomotion but did increase cFos in the BLA. **A)** Graphs showing the effect of optogenetic inhibition of PV+ cells in the BLA on locomotion (left) and anxiety-like behavior (right). Optogenetic inhibition of PV+ cells did not affect distance traveled ($p$ = 0.72) or the time spent in the center of the open field ($p$ = 0.19) when compared to light controls. For light control n = 7, for PV Halo n = 6. **B)** Graph showing the effects of optogenetic inactivation on cFos labeling within the BLA, following 15 minutes of unilateral inhibition in the home cage. Immunohistochemical labeling for c-Fos revealed that unilateral optogenetic inactivation of PV+ cells increased c-Fos positive cells in the inhibited side of the brain ($p$ = 0.04) when compared to the uninhibited side (n = 5 per group). A two-tailed paired student t.test was used to assess differences in c-Fos labeling, for all other analysis a two-tailed unpaired student t.tests were used. For all graphs the mean and SEM are presented. * = $p < 0.05$, ** = $p < 0.01$, *** = $p < 0.001$.
